# Half as high for twice as long: patriarchy and the age-structure of human fertility

**DOI:** 10.1101/2025.05.19.654849

**Authors:** Matthew C. Nitschke, Kristen Hawkes, Peter S. Kim

## Abstract

Humans are distinguished from our closest living relatives, other great apes, by our extended postmenopausal longevity, later first births, and shorter birth intervals. Those features likely evolved as ancestral grandmothers’ foraging subsidized dependants in habitats lacking foods that youngsters could manage for themselves. If so, as female post-fertile years increased, older years increased in males too. Those still-fertile old males in the paternity competition pushed the average male fertility rate below the female average. As R. A. Fisher explained, Mendelian inheritance requires equal contributions from both sexes to descendant gene pools. Higher fertility in one sex makes tendencies to overproduce it pass to more grandchildren, equalising averages in descendant generations. Yet lower average number of offspring per year in men persists, their fertility lasting decades longer than women’s. Here, we present a simple mathematical model to investigate this phenomenon. We show that a male-biased sex ratio in the fertile ages is consistent with an offspring sex ratio of 1:1. We show that if male fertile careers are twice as long as those of females, the extended male fertility soon balances the higher female average, maintaining Fisher’s equilibrium offspring sex ratio under human life history conditions.

**Significance Statement:** Why are offspring sex ratios usually near even in humans when the fertile careers of women and men differ so much? Fisher’s principle explains why parents are usually expected to produce equal offspring sex ratios. Since every sexually produced grandchild has both a mother and father, the average number of grandchildren expected through offspring of either sex depends on the sex ratio of the fertile pool. If that sex ratio is biased, the average must be higher for the rarer sex. Then any tendency to overproduce the sex with higher average fertility will spread, reducing its rarity until expected averages through sons and daughters are equal. Human life history complicates this simple logic. Female fertility ends in mid-life whereas male fertility does not, creating a male-biased fertile-age population in which average number of offspring per year is lower in males than in females. We show that sex differences in fertility duration can maintain Fisher’s equilibrium offspring sex ratio by altering Reproductive Value in overlapping generations, providing a foundation for persistent patriarchy.

## Introduction

Fisher’s principle follows from a simple feature of sexual reproduction in which every offspring has one mother and one father, so each sex must supply half the autosomal ancestry to future generations. In populations with overlapping generations, one sex may contribute fewer offspring per year but continue producing offspring over a longer fertile span. Thus, selection is affected by the rate males and females produce offspring, and also when they produce them.

In humans, female fertility ends before menopause while male fertility continues much longer. Fertile-age males can therefore outnumber fertile-age females, making daughters appear to provide the higher average number of grandchildren. Yet human offspring sex ratios remain nearly equal. Resolving this apparent contradiction requires following gene copies across overlapping generations and asking how links with social behavior change the timing of maternity and paternity. Hamilton’s inclusive fitness argument extends the gene’s-eye view to behaviors that affect others’ survival or fertility. A gene linked to behavior can be favored when the fitness benefit it provides to others sharing a copy of the gene exceeds the fitness cost to the actor, as expressed by Hamilton’s rule, *rb > c* [1]. With sexual reproduction, diploid offspring get half their genes from each parent, making the chance a rare gene identical by descent is in an offspring (*r* = 0.5) twice the chance it is in a grandoffspring (*r* = 0.25).

Here, we place this idea in a human social-behavioral setting. The evolution of our grandmothering life history [2–4] can explain why human populations are biased toward males in the fertile ages [5–9]. With that male bias, unusual for a mammal, mate-guarding wins more paternities than multiple mating [10–15]. If deference to older males’ claims raises average paternal ages above average maternal ages, then generational steps from fathers put more genes from previous generations into generations produced by younger mothers.

To clarify this and its consequences, we consider an extreme example. Assume that the fertile ages of males are twice as long as those of females. The model builds on Grafen’s age and sex-structured treatment of reproductive value in diploid populations with overlapping generations [16]. Grafen traced a gene’s ancestry from a distant future generation as it jumps back into a mother or a father at each generational step, showing that Reproductive Value totaled across females and males depends on average maternal and paternal ages. Subsequent work has extended this logic to other inheritance systems, including X-linkage and haplodiploidy [17], and more generally to juvenile and total Reproductive Values under arbitrary genetic systems [18]. These studies establish the formal Reproductive Value framework, but *they do not ask how particular social and life-history conditions generate differences in the age distribution of maternity and paternity*. Here, we apply this reasoning to the human case, where the sex ratio of fertile adults is male-biased. That makes the average rate of offspring production per year lower in fertile males than it is in fertile females while male fertility lasts decades longer. Our aim is to show how differences in paternity, fertility duration, and parental age can alter sex-specific Reproductive Values (defined by Fisher as expected future fertility) while remaining consistent with Fisher’s principle about equal offspring sex ratios.

### Background

It has long been recognized that sex differences in adult mortality need not alter Fisher’s argument about equal offspring sex ratios. If adult sons are twice as likely to die as adult daughters, then the sons that survive have double the paternity chances (see Figure 1(a)).Here, we investigate whether the difference in fertile spans between the sexes in humans might balance in the same way (Figure 1(b)) [19],consistent with the general result derived by Grafen.

**Figure 1.**
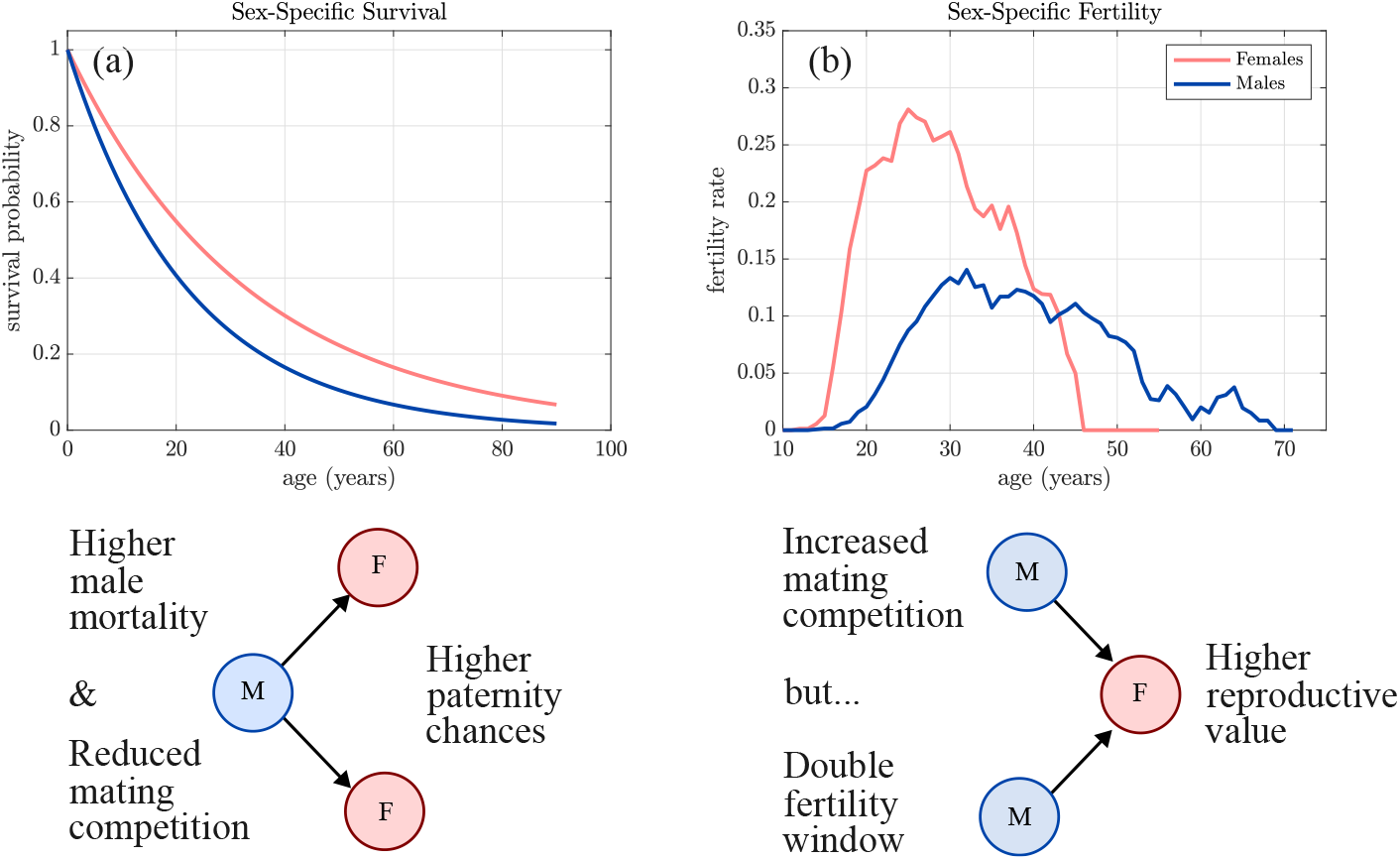
Sex-biased fertile populations and reproductive consequences. Illustrations depict two demographic scenarios. (a) When mortality is higher in adult males, that leaves fewer competitors and so more paternities for survivors. (b) In a male-biased population, intensified male competition reduces males’ short-term reproductive success, but their longer fertility window increases the lifetime reproductive value of sons. Both scenarios reflect frequency-dependent selection consistent with Fisher’s principle: reproductive value adjusts to maintain equal genetic contribution from males and females to subsequent generations, even in the presence of age-specific mortality or fertility asymmetries.

Darwin had recognized the one-mother-one-father axiom for sexual reproduction [20]; however, it was Fisher [21], combining natural selection with Mendelian inheritance, who saw that each sex contributes half the autosomal genes to future generations which usually pushes natural selection to equal expenditure on sons and daughters. Hamilton later summarized the same logic, noting that if male births are less common, “a newborn male then has better mating prospects than a newborn female” [22].This logic, generally referred to as the “Fisher Principle”, makes the average fertility of the more abundant sex lower than the average of the rarer one, so tendencies to bias offspring toward the rarer sex occur in more grandchildren. That rare sex advantage, first modelled by Karl Düsing [23], is widely considered the first identification of frequency-dependent selection [24].

More generally, rare sex advantage explains why there are so many males, even when–as in most primates–the number of babies depends only on the number of fertile females. If males are rare and one or a few males can fertilize all the females almost simultaneously, the average success of rare males must be much higher than the female average. Tendencies to bias offspring toward the rare males would be more common in the grandoffspring generation, reducing males’ rarity. An equilibrium is reached when the average reproductive success of the two sexes is equal, demonstrating an evolutionarily stable strategy (ESS) [25].

We explore the possibility that inclusive fitness consequences of deference to older males’ mate-guarding could balance a sex difference in average fertilities with an extreme simplification. Assume that average male fertility is half the female average but extends over twice as many fertile ages. Then, although sons average half the offspring output of daughters, their fertile careers are twice as long, so they put more genes into future generations than daughters do.

This extreme example shows that the number of accumulated replicas through sons begins to catch up to the number through daughters. Since mothers bear the offspring, generation length depends only on maternal ages; fathers’ ages are irrelevant to generation length. However, fathers’ ages do matter for selection because traits that increase success in paternity competition accumulate in subsequent generations. While a hypothetical offspring gene gets only half the replicas through sons as through daughters each generation at time *t*, it has double the chance of putting copies in the next generation at time *t* + 1. A gene is being favoured by natural selection if the aggregate of its replicas forms an increasing fraction of the total gene pool [26].Population genetic models of senescence have shown that the selection pressure on survival is a function of Reproductive Value at any age, weighted by the probability of surviving to that age [27, 28].However, these models make use of Fisher’s explicit definition of Reproductive Value (expected future fertilities) that leaves out the inclusive contributions of still-productive post-fertile females [19]. Fisher showed that when age-specific fertility and mortality remain constant and migration is negligible, populations attain a stable age structure in which one can compute the direct contribution from each age class to future gene pools by combining survival and fertility rates across generations. Although he briefly acknowledged “there will also, no doubt, be indirect effects in cases in which an animal favours or impedes the survival or reproduction of its relatives…as in mankind a mother past bearing age may greatly promote the reproduction of her children” [21], he surmised that “such indirect effects will in very many cases be unimportant”. By ignoring those indirect effects, he could take advantage of the renewal equation, which attends only to age-specific fertilities and mortalities. It was the importance of including both direct and indirect effects that Hamilton would later elaborate and formalize as inclusive fitness [1].

### The demographic basis of the model

Careful collection and analysis of demographic data for modern human foragers living under mortality regimes, more like those that prevailed through most of human experience, show that post-fertile women are not rare [29–32]. Instead, females that survive to adulthood are highly likely to live beyond their fertility window. Human age structures include substantial fractions of post-fertile women. Thus, although postmenopausal women do not produce offspring, they must indeed be characterized by a genetic reproductive value not too different from that of males [33]. Human age structures also include substantial fractions of old men who are still fertile. Some out-compete younger men for paternities. The strong male bias of the sex ratio in the fertile ages which makes that possible distinguishes humans not only from the other great apes [34], but from most other mammals. This is illustrated for several modern hunter-gatherer populations in Figure 2. These figures motivate the simplified model below. Female fertility is concentrated into a shorter window, whereas male fertility extends longer. This lowers average male offspring production by increasing the number of competing fertile males, while allowing genetic contributions through sons to accumulate over a longer fertility window.

**Figure 2.**
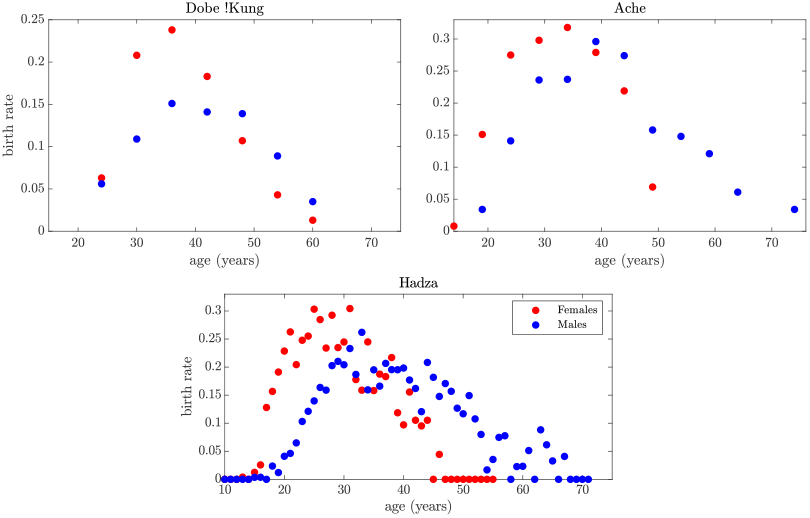
Age-specific fertility for women and men in three modern hunter-gatherer societies. Scatter plots show empirical estimates of age-specific fertility for women (red) and men (blue) among the Dobe !Kung [29] (top left), Aché [30] (top right), and Hadza [32] (bottom center). Female fertility data exhibit a narrower reproductive span, typically peaking in the late twenties and declining steeply by age 45. Male fertility spans a broader age range and averages decline more gradually, particularly in the Aché and Hadza. All data are derived from published demographic reports and are plotted as observed. These distributions illustrate persistent sex differences in fertility timing and duration, which underpin the sex-specific structure of Reproductive Values in later analyses.

In humans, as is common in mammals generally [35, 36], mortality is higher in males. As noted above, higher male mortality in early life has been widely understood to imply that the slight male bias in human birth sex ratios regularly found in large samples is part of an adjustment from an unexplained higher male bias at conception with early male-biased mortality allowing replacements that result in equal offspring sex ratios as predicted by Fisher’s account. However, a recent empirical study showed sex ratios near equal at conception, contradicting the claim that the sex ratio at conception is male-biased [37, 38].Our concern here is what happens in adult ages, where higher levels of adult mortality in males are sufficiently remarked in humans to pose a “health-survival paradox” [19, 39].

Higher adult mortality in males does raise paternity opportunities for the survivors. But stopping there to account for near equal offspring sex ratios in humans misses a distinctive and consequential feature of our fertile age structure: Old men only gain paternities by outcompeting younger men for claims on younger women.

## Results

We ask whether, given the sexual asymmetry in human fertile ages, males producing offspring at a rate lower than the female mean can still satisfy Fisher’s equal offspring sex ratio principle. To investigate this, we begin with a simplified age-and sex-structured model in which males reproduce on average half the average female rate but remain fertile for twice as long. We show that this difference in fertility timing is consistent with an equal offspring sex ratio. We then examine the same demographic asymmetry in Reproductive Value terms by tracing gene copies back through sequential conceptions from a distant future generation. This shows how extended male fertility alters the distribution of ancestry across male and female age classes.

### Balanced offspring sex ratio

Consider a population of females, *F*, and males, *M*, divided into *ω* age groups, with the number in each group at time *t* given by the vector

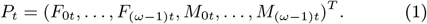

We define a genetic projection matrix **K** which tracks genetic contributions from births by assigning half of each offspring’s ancestry to the mother and half to the father. Furthermore, we assume a stationary population with an equal offspring sex ratio, thus **K** tracks the long-term transmission of gene copies through daughters and sons.

To determine the contribution of one newborn female to subsequent descendants at time step *n*, we start with the initial condition

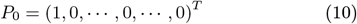

and calculate **K**^*n*^*P*_0_, with the contribution of one newborn female to subsequent descendants. By construction, the population will approach a stable age distribution. Thus,

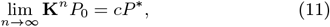

for some constant *c*. Note that over her lifetime, one newborn female will have a total of one female descendant and one male descendant. Thus, she will replace herself (we assumed a stationary population). Similarly, sons replace themselves under an equal offspring sex ratio, thus one newborn male will have a total of one female and one male descendant. Since the property of self-replacement is invariant, i.e., future descendants will replace themselves, both initial conditions of one newborn female and one newborn male will approach the stable age distribution

Thus, whether you start with one newborn male or one newborn female, the long-term output through offspring converges to the same stable age distribution. This means that each will have the same long-term contribution to future offspring, which implies that an equal offspring sex ratio is the optimal evolutionary strategy. Hence, an offspring sex ratio of 1:1 is consistent with Fisher’s principle. An illustration of this is shown in Figure 3 where all males in the fertile ages have an equal chance at a paternity. Our numerical simulation includes an equally-weighted carrying capacity under which individuals are removed from the population with equal probability from all age classes.

**Figure 3.**
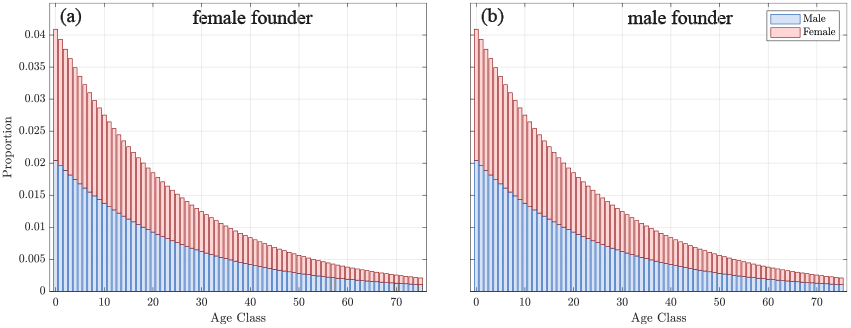
Stable age distributions produced by the two-sex 2*ω* × 2*ω* Leslie matrix model under sex-specific founder conditions. Each subplot displays the normalised age structure at *t* = 500 following initialisation with a single newborn individual of either a female founder (a) or a male founder (b). Stacked bars represent the proportion of females and males in each age class. For simplicity, we set the sex-specific survival rates as *s*_*f*_ = *s*_*m*_ = 0.98, *q*_*m,i*_ = 1, and assume constant per capita male and female fecundity in accordance to our half-as-high-for-twice-as-long example with *f*_*f,i*_ = 0.2 for *i* ∈ {15, …, 45} and *f*_*m,i*_ = 0.1 for *i* ∈ {15, …, 75}. Note that both males and females will have the same long-term contribution to future offspring, implying an equal offspring sex ratio is the optimal strategy.

### Older Fathers Balance Lower Male Fertility

Next, we consider the trajectory of a gene backward in time and show that Reproductive Values of all males and of all females depend on the age of an average new father and new mother. Following a thought experiment suggest by Alan Grafen [16], we look back at a gene’s history from far in the future as it jumps back into a mother or father in each generation. He pointed out that although non-overlapping generations ensure equal Reproductive Value of females and males in each generation, overlapping generations will cause differences to arise between them. Consider an idealized model in which females are fertile for one fertile age class and reproduce with probability 1. This allows for mathematical simplicity and greater clarity. The overall age structure for this model is illustrated in Figure 4.Within the fertile age classes, we assume males are fertile for two generations, corresponding to ages approximately 15 to 75, exactly double that of females who are fertile from the age of 15 to 45.

**Figure 4.**
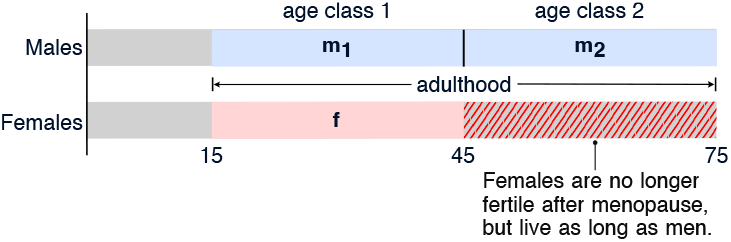
Age classes for males and females. In our simplified example, we consider one age class for females, corresponding to ages of female fertility, approximately ages 15 to 45. Males have two fertile age classes, *m*_1_ and *m*_2_, corresponding to young ages 15 to 45 and old ages up to age 75.

Let *w*_1_ denote the probability that an ancestral path moving through a parent passes through a young father, *m*_1_, and let *w*_2_ = 0.5 *™ w*_1_ be the probability it passes through an old father, *m*_2_. The remaining probability, 0.5, corresponds to transmission through the mother. Thus, conditional on the path passing through a father, the probabilities of young and old fathers are 2*w*_1_ and 2*w*_2_, respectively (see supplementary information). We also assume that there is an equal offspring sex ratio and the population is stationary, so in each time step a female will give birth to one son and one daughter.

Figure 5 illustrates the path a single gene may take starting from a distant time in the future *T* back to time *T ™* 1. The ancestral path passing through a female or male of age 1 has three possible backward transitions: to a female parent at the previous time step with probability 0.5, to an age-1 male parent with probability *w*_1_, and to an age-2 male parent with probability *w*_2_. Since *w*_1_ + *w*_2_ = 0.5, these probabilities sum to one. A male of age 2 transitions back to his younger self, a male of age 1 with probability 1.

**Figure 5.**
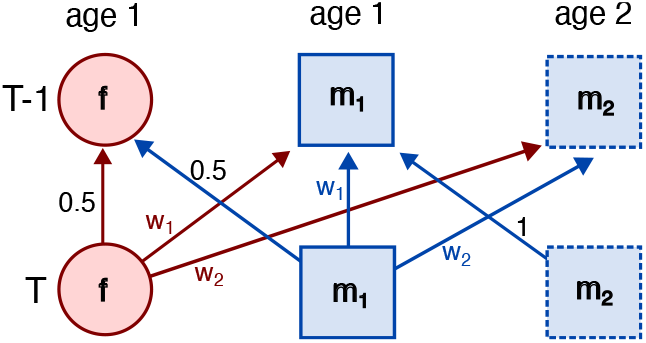
Ancestral pathways in a backward-time Markov model. Diagram showing the probabilistic paths of a single autosomal gene tracing back one generation from a future time *T*. Probabilities on arrows reflect the frequency with which a gene is expected to pass through individuals of each sex and age, supporting the inference of sex-specific contributions to offspring over time.

The equilibrium distribution over these three states *f*, *m*_1_ and *m*_2_ is

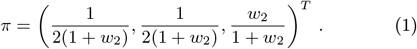

This is used to determine the total lifetime Reproductive Value summed over all ages, *a*, for females and males respectively:

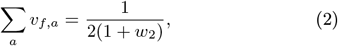

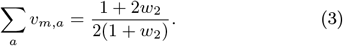

Thus, as long as age class *m*_2_ has a non-zero chance of winning some paternities, *w*_2_ *>* 0, the total reproductive value of males is greater than that of females:

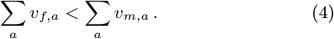

Using the data illustrated in Figure 2, we approximate the distribution of male paternities as 85% in the younger male age class (~15 - 45 yrs.) and 15% in the older male age class (~46 - 85 yrs.). Since paternal ancestry accounts for one half of autosomal transmission, this corresponds to *w*_1_ = 0.425 and *w*_2_ = 0.075. Thus, at a reproductive transition, 50% of autosomal ancestry paths pass through *f*, 42.5% pass through *m*_1_ and 7.5% pass through *m*_2_. Therefore, substituting *w*_2_ = 0.075 into Eqs. 2 and 3, the total reproductive value for males and females is

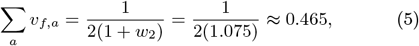

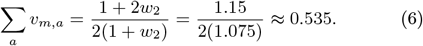

Thus, even though the fertility of males is approximately half that of females, their lifetime Reproductive Value is higher since older fathers put more genes from previous generations into generations through younger mothers. Thus, the number of aggregate replicas through sons begins to catch up to the number through daughters. This effectively balances sex differences in the average fertilities of those in the fertile ages.

## Discussion

It has long been recognised that sex differences in adult mortality do not necessarily violate Fisher’s argument for equal parental investment in sons and daughters. The question in humans is whether a male-biased fertile-age sex ratio, which lowers average male fertilities, should favour daughters instead. Our results provide a demographic illustration of Grafen’s [16]result that differences in age at offspring production can generate differences in Fisherian Reproductive Value while preserving Fisher’s equal investment condition for offspring.

Although average offspring production is lower in fertile males than in fertile females, men remain fertile through adulthood while women’s fertility ends in mid-life. Men’s extended fertility enables them to accumulate genetic contributions over a longer period, thereby compensating for their lower average fertility. We explored this scenario using two mathematical models, showing that if male fertile careers are twice as long as those of females, the corresponding Reproductive Value (defined as future fertility) of males is larger. Thus, extended male fertility balances the sex differences in average rate of offspring production, preserving Fisher’s offspring sex ratio equilibrium for human life histories. Even though we illustrate this with the ‘half-as-high-for-twice-as-long’ scenario, an example chosen for simplicity, the modelling framework applies to any sex difference in fertility duration or fertility rates.

Our models suggest a plausible evolutionary scenario in which sex differences in the duration of fertility are consistent with a balanced offspring sex ratio and equal long-term genetic contributions from males and females. The relevant human asymmetry is not only physiological. If ancestral grandmothers’ foraging subsidies propelled the evolution of our postmenopausal longevity and shortened birth intervals, the resulting distinctively human age-structure of fertility puts more males in competition for younger females. With more competitors, mate guarding wins more paternities. Deference to proprietary claims of older males is amplified by reputations, making competition for standing an incessant feature. We do not explicitly model male hierarchies or deference implications here (but see [40, 41]).However, our modelling shows that even without these assumptions, the demographic consequences of human fertile-age structure and postmenopausal longevity amplify sex-specific differences in Fisher’s reproductive value. By extending male fertility beyond female fertility, this structure likely set the foundation for persistent human patriarchy.

## Materials and methods

We used both a modified Leslie matrix and Markov chain model to obtain our results.

### Leslie Matrix model

Starting with the population of males and females, the change in population structure from time *t* to *t* + 1 can be represented by a projection of a two-sex Leslie-type matrix

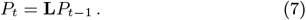

The matrix **L** has the standard Leslie structure. Survival elements *s*_*f,i*_ and *s*_*m,i*_ move females and males from age class *i* to *i* + 1, while reproductive rows place newborn daughters and sons into age class zero. We assume a stationary population and write *l*_*f,i*_ and *l*_*m,i*_ for survival to age class *i*, with *l*_*f*,0_ = *l*_*m*,0_ = 1. Female fecundity is scaled so that a newborn female exactly replaces herself over her lifetime

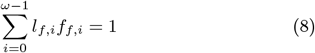

and we assume a balanced offspring sex ratio. Thus, the fecundity terms in rows 1 and *ω* + 1 of **L** are symmetric (see Supplementary Information).

Let

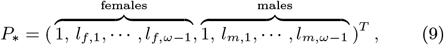

where *l*_*f*,0_ = 1, *l*_*f,i*_ = *s*_*f*,0_*s*_*f*,1_ *… s*_*f,i™*1_, *l*_*m*,0_ = 1 and *l*_*m,i*_ = *s*_*m*,0_*s*_*m*,1_ *s*_*m,i™*1_ for *i ≥* 1. *P*_*∗*_ is the stable age distribution, so is the eigenvector of **L** corresponding to the eigenvalue 1. Hence any age distribution of the population will be proportional to *P*_*∗*_.

Male paternity success may vary by age. Let *q*_*m,i*_ *≥* 0 denote the age-specific paternity weight of males in age class *i*. At the stable age distribution, the weighted male competition pool is

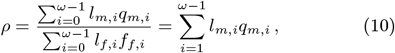

where we have used the female normalisation 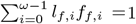. We use this to define the normalised age-specific paternial contribution as

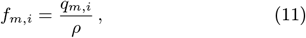

so that

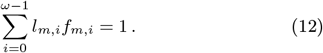

Inserting this into the two-sex Leslie matrix to account for paternities and assuming an offspring sex ratio of 1:1, the modified Leslie matrix, **K**, that tracks how genetic contributions pass from one generation to the next is

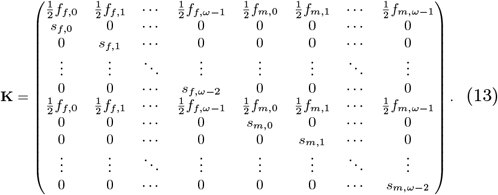

The first and (*ω*+1)th rows of **K** are identical because we assume an equal offspring sex ratio. Each birth term is weighted by 1*/*2, reflecting autosomal inheritance from the mother or father, while the subdiagonal terms describe sex-specific survival. With the fecundities normalised as above, **K** has the same stable age distribution as **L**.

Starting from a single newborn female,

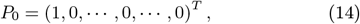

the descendant contribution after *n* time steps is **K**^*n*^*P*_0_. Since the population is stationary and both female and male fecundities are normalised to self-replacement, founder lineages of either sex converge to the same stable age distribution:

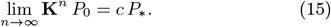

The scaling constant is

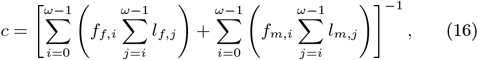

the reciprocal of the total lifetime descendant output generated by the founder vector.

### Markov Chain Model

The backward-time transitions in Figure 5define the columnstochastic Markov matrix

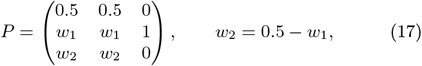

where the states are a female parent, a young male parent, and an older male parent, respectively. Solving *Pπ* = *π* gives

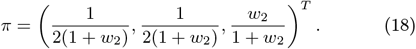

The first two components correspond to parent–offspring transitions, whereas the final component is the within-individual aging transition from an older male to his younger self. Hence the probability that an ancestral path moves between individuals is

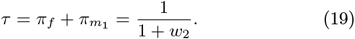

Let *u*_*s,a*_ be the probability that the ancestral path moves through a parent of sex *s* and age class *a*. In the stable population,

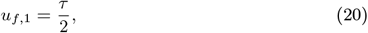

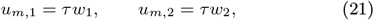

with all other age classes contributing zero in this idealised example.

The reproductive value of a newborn of sex *s* is obtained by summing over future ancestral transitions,

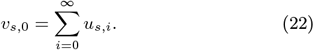

Therefore,

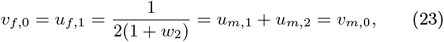

so newborn females and newborn males have equal reproductive value, as required by Fisher’s principle.

Total reproductive value over reproductive ages differs because males occupy two reproductive age classes:

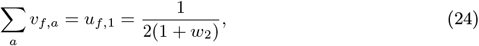

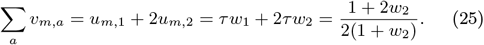

## Data, Materials, and Software Availability

The authors confirm that all empirical data used in this study derive from previously published demographic sources, as cited in the article. The processed data and all MATLAB code used to generate the analyses and figures are available at https://github.com/mcnitschke/half-as-high. A permanent archived version is available at https://doi.org/10.5281/zenodo.17836695.

## Acknowledgments

The authors gratefully acknowledge support for this work through the Australian Research Council Future Fellowship FT200100190 (PSK).

## Author contributions

Matthew C. Nitschke: Methodology, Formal analysis, Investigation, Writing – original draft, Writing – review & editing, Visualization. Kristen Hawkes: Conceptualization, Writing – original draft, Writing – review & editing. Peter S. Kim: Conceptualization, Methodology, Formal analysis, Writing –review & editing.

