## Supplementary Information for "Half as high for twice as long: patriarchy and the age-structure of human fertility"

### 1 Leslie matrix model

In this section, we consider a diploid population in discrete time and introduce a two-sex Leslie matrix model. Using this model, we show that an equal offspring sex ratio is the optimal evolutionary strategy even when the age-specific fertility difference between the sexes is male biased (males have a longer fertility window).

Table 1 provides a summary of parameters and population variables used in our model.

| Variable | Description |
|---|---|
| $F_i$ | Number of females in the age class $i$ |
| $M_i$ | Number of males in the age class $i$ |
| $s_{f,i}$ | Fraction of females who survive from age class $i$ to $i+1$ |
| $s_{m,i}$ | Fraction of males who survive from age class $i$ to $i+1$ |
| $l_{f,i}$ | Probability of females surviving to age $i$ |
| $l_{m,i}$ | Probability of males surviving to age $i$ |
| $f_{f,i}$ | Per capita female fecundity of age class $i$ |
| $f_{m,i}$ | Per capita male fecundity of age class $i$ |
| $q_{m,i}$ | Age-specific paternity weight of males in age class $i$ |

Table 1: Parameters and Population Variables

Consider a population of females and males divided into $\omega$ age groups, with the number in each group at time $t$ given by the vector

$$P_t = (F_{0t}, \dots, F_{(\omega-1)t}, M_{0t}, \dots, M_{(\omega-1)t})^T, \tag{1}$$

where superscript $T$ denotes the transpose. The change in population structure from time $t$ to $t+1$ is then represented by the equation

$$P_t = LP_{t-1}. \tag{2}$$

where $L$ is a two-sex Leslie matrix in which $f_{f,i}$ are the fecundities of mothers, $s_{f,i}$ are the fractions of females, and $s_{m,i}$ the fractions of males who survive from age class $i$ to $i+1$. A more explicit description of this model is shown in equation (3).

$$L = \begin{pmatrix} f_{f,0} & f_{f,1} & \cdots & f_{f,\omega-2} & f_{f,\omega-1} & 0 & 0 & \cdots & 0 \\ s_{f,0} & 0 & \cdots & 0 & 0 & 0 & 0 & \cdots & 0 \\ 0 & s_{f,1} & \cdots & 0 & 0 & 0 & 0 & \cdots & 0 \\ \vdots & \vdots & \ddots & \vdots & \vdots & \vdots & \vdots & \ddots & \vdots \\ 0 & 0 & \cdots & s_{f,\omega-2} & 0 & 0 & 0 & \cdots & 0 \\ f_{f,0} & f_{f,1} & \cdots & f_{f,\omega-2} & f_{f,\omega-1} & 0 & 0 & \cdots & 0 \\ 0 & 0 & \cdots & 0 & 0 & s_{m,0} & 0 & \cdots & 0 \\ 0 & 0 & \cdots & 0 & 0 & 0 & s_{m,1} & \cdots & 0 \\ \vdots & \vdots & \ddots & \vdots & \vdots & \vdots & \vdots & \ddots & \vdots \\ 0 & 0 & \cdots & 0 & 0 & 0 & 0 & \cdots & s_{m,\omega-2} \end{pmatrix}. \tag{3}$$

Note that the $\omega \times \omega$ matrix in the upper left corner of $L$ is the standard Leslie matrix model for a female population. The lower $\omega$ rows of the matrix account for the survival probabilities and births of new males from the female population. We assume a stationary population with no growth, so $\lambda = 1$. Let $l_{f,i}$ be the probability a female survives multiple age classes up through age class $i$, then $l_{f,i} = s_{f,0} \cdot s_{f,1} \cdots s_{f,i-1}$ for $i \geq 1$. Each female is born with certainty, so $l_{f,0} = 1$. Thus, the female parameters satisfy the Euler-Lotka equation

$$\sum_{i=0}^{\omega-1} l_{f,i} f_{f,i} = 1. \tag{4}$$

This means that the expected lifetime reproductive success of a newborn individual is one and each individual exactly replaces itself. Additionally, we assume a balanced offspring sex ratio so that sons and daughters are produced in equal numbers from the same fertility distribution. Thus, the fecundity terms in rows 1 and $\omega + 1$ of $L$ are symmetric.

Let

$$P^* = (\underbrace{1, l_{f,1}, \ldots, l_{f,\omega-1}}_{\text{females}}, \underbrace{1, l_{m,1}, \ldots, l_{m,\omega-1}}_{\text{males}})^T. \tag{5}$$

where $l_{f,0} = 1$, $l_{f,i} = s_{f,0} s_{f,1} \cdots s_{f,i-1}$, $l_{m,0} = 1$ and $l_{m,i} = s_{m,0} s_{m,1} \cdots s_{m,i-1}$ for $i \geq 1$. $P^*$ is the stable age distribution, that is, it is an eigenvector of the two-sex Leslie matrix corresponding to the eigenvalue 1. Hence, any age distribution of the population will be proportional to $P^*$.

For simplicity, we assume all surviving males are able to compete for paternities, but allow their expected paternity success to vary by age. Let $q_{m,i} \geq 0$ denote the age-specific paternity weight of males in age class $i$. In addition, assume the population has reached a stable age distribution. This means that the age-specific fertility is constant, while male paternity success is distributed across age classes according to the weights $q_{m,i}$. Thus, we can define $\rho$, the weighted male competition pool relative to females who are able to conceive in the next time step as:

$$\rho = \frac{\sum_{i=0}^{\omega-1} l_{m,i} q_{m,i}}{\sum_{i=0}^{\omega-1} l_{f,i} f_{f,i}}. \tag{6}$$

The denominator of this ratio is 1 from Eq. 3, so it becomes

$$\rho = \sum_{i=0}^{\omega-1} l_{m,i} q_{m,i}. \tag{7}$$

We wish to construct a modified Leslie matrix to track how many descendants come from males and females. Using the weighted male competition pool defined above, we allow male fecundity to vary by age. In particular, the expected paternity contribution of a male in age class $i$ is proportional to $q_{m,i}$ and is normalised by the total weighted male competition pool, $\rho$. Thus, the age-specific fecundity of males is

$$f_{m,i} = \frac{q_{m,i}}{\rho}. \tag{8}$$

By construction,

$$\sum_{i=0}^{\omega-1} l_{m,i} f_{m,i} = 1 \tag{9}$$

so the total paternity output is normalised across all surviving male age classes.

Inserting this into the two-sex Leslie matrix above to account for paternities and assuming an offspring sex ratio of 1:1, the modified Leslie matrix, $K$, that calculates how genetic contributions pass from one generation to the next is

$$K = \begin{pmatrix} \frac{1}{2}f_{f,0} & \frac{1}{2}f_{f,1} & \cdots & \frac{1}{2}f_{f,\omega-1} & \frac{1}{2}f_{m,0} & \frac{1}{2}f_{m,1} & \cdots & \frac{1}{2}f_{m,\omega-1} \\ s_{f,0} & 0 & \cdots & 0 & 0 & 0 & \cdots & 0 \\ 0 & s_{f,1} & \cdots & 0 & 0 & 0 & \cdots & 0 \\ \vdots & \vdots & \ddots & \vdots & \vdots & \vdots & \ddots & \vdots \\ 0 & 0 & \cdots & s_{f,\omega-2} & 0 & 0 & \cdots & 0 \\ \frac{1}{2}f_{f,0} & \frac{1}{2}f_{f,1} & \cdots & \frac{1}{2}f_{f,\omega-1} & \frac{1}{2}f_{m,0} & \frac{1}{2}f_{m,1} & \cdots & \frac{1}{2}f_{m,\omega-1} \\ 0 & 0 & \cdots & 0 & s_{m,0} & 0 & \cdots & 0 \\ 0 & 0 & \cdots & 0 & 0 & s_{m,1} & \cdots & 0 \\ \vdots & \vdots & \ddots & \vdots & \vdots & \vdots & \ddots & \vdots \\ 0 & 0 & \cdots & 0 & 0 & 0 & \cdots & s_{m,\omega-2} \end{pmatrix}. \tag{10}$$

The first $\omega$ elements of the first row represent the number of daughters produced by females in the current time step multiplied by a factor of 1/2 as descendants carry half the genes of their mothers. The last $\omega$ elements of the first row are the number of daughters produced by males in the current time step. These elements are also multiplied by 1/2 since descendants carry half the genes of their fathers. We have assumed an equal offspring sex ratio, thus the $(\omega + 1)$th row, which corresponds to the number of sons produced in the current time step, is the same as the first row. $K$ has the same stable age distribution as $L$, that is, they have the same leading eigenvalue of 1.

To determine the contribution of one newborn female to subsequent descendants at time step $n$, we start with the initial condition

$$P_0 = (1, 0, \cdots, 0, \cdots, 0)^T \tag{11}$$

and calculate $K^n P_0$, with the contribution of one newborn female to subsequent descendants. By construction, the population will approach a stable age distribution. Thus,

$$\lim_{n\to\infty} K^n P_0 = cP^*, \tag{12}$$

for some constant $c$. Note that over her lifetime, one newborn female will have a total of one female descendant and one male descendant. Thus, she will replace herself (we assumed a stationary population). Also, by the way we defined fecundity in terms of the sex ratio, over his lifetime, one newborn male will also have a total of one female and one male descendant. Since the property of self-replacement is invariant, i.e., future descendants will also replace themselves, both initial conditions of one newborn female and one newborn male will approach the stable age distribution $cP^*$ with

$$c = \left[ \sum_{i=0}^{\omega-1} f_{f,i} \left( \sum_{j=i}^{\omega-1} l_{f,j} \right) + \sum_{i=0}^{\omega-1} f_{m,i} \left( \sum_{j=i}^{\omega-1} l_{m,j} \right) \right]^{-1}, \tag{13}$$

where $c$ is the reciprocal of the total lifetime number of descendants produced by the initial population vector $P_0$.

### 2 Markov Chain Model

Table 2 provides a summary of parameters used in the Markov Chain model.

| Variable | Description |
|---|---|
| $f$ | Female reproductive age class |
| $m_1$ | Younger reproductive age class |
| $m_2$ | Older reproductive age class |
| $p_{t,s,a}$ | Probability a new parent at time $t$ is sex $s$ and age $a$. |
| $u_{t,s,a}$ | Probability a transmitting ancestor at time $t$ is sex $s$ and age $a$. |
| $\tau_t$ | Fraction of ancestral paths at time $t$ that move between individuals. |
| $v_{t,s,a}$ | Total ancestral-path probability for sex $s$, age $a$, summed over future ages. |
| $u_{t+i,s,a+i}$ | Transmission probability $i$ time steps later, when age $a$ has become $a+i$. |

Table 2: Notation used in the Markov chain model.

Following [1], we define the probability that a new parent is of age $a \in \{1, 2, 3, \ldots\}$ and sex $s \in \{m, f\}$ at time $t$ as $p_{t,s,a}$. Since we assume a diploid dioecious population and all females are fertile and males eligible for a paternity with only two age classes,

$$p_{t,f,1} = 1, \tag{14}$$

$$p_{t,m,1} = 2w_1, \quad p_{t,m,2} = 2w_2, \quad w_1 + w_2 = 0.5. \tag{15}$$

Additionally, we define $\tau_t$ as the chance an ancestral path move between individuals at time $t$. In other words, this is the fraction of paths at time $t$ that transition from a parent to a newborn. Since we assume a stable age distribution, $\tau_t$ is the same for all $t$.

The ancestral path passing through a female or male of age 1 has three possible backward transitions: to a female parent at the previous time step with probability 0.5, to an age-1 male parent with probability $w_1$, and to an age-2 male parent with probability $w_2$. Since $w_1 + w_2 = 0.5$, these probabilities sum to one. A male of age 2 transitions back to his younger self, a male of age 1 with probability 1.

The probabilities define a transition matrix for the Markov chain

$$P = \begin{pmatrix} 0.5 & 0.5 & 0 \\ w_1 & w_1 & 1 \\ w_2 & w_2 & 0 \end{pmatrix}, \qquad w_2 = 0.5 - w_1, \tag{16}$$

where each column corresponds to the transition for a gene copy currently carried by a female, an age-1 male, or an age-2 male in generation $T$. This allows us to determine the long-term stationary distribution of the system. Let $\pi$ denote the equilibrium distribution over the three states $f$, $m_1$, and $m_2$, then

$$\pi = \left(\frac{1}{2(1+w_2)}, \frac{1}{2(1+w_2)}, \frac{w_2}{1+w_2}\right)^T. \tag{17}$$

This means that at equilibrium, the fraction of ancestral paths passing through females is $[2(1+w_2)]^{-1}$, the same fraction passes through age-1 males, and the fraction passing through age-2 males is $w_2(1+w_2)^{-1}$. This means that the fraction of paths involving a transition between individuals is the sum of the female and age-1 components. The remaining fraction corresponds to the within-individual transition from an age-2 male back to his age-1 self. Hence,

$$\tau = \frac{1}{1+w_2}. \tag{18}$$

Next, define $u_{t,s,a}$ as the probability that an autosomal gene's unique ancestral path passes to an offspring from an individual of a given age and sex at time $t$. In order for this to happen, the ancestral path must move between individuals (not due to aging) and additionally, there must be a new available parent of the correct age, $a$, and sex, $s$, at time $t$. Thus, since $\tau$ is the chance that the path moves between individuals and $p_{t,s,a}$ is the probability that a specific parent is available, we can define

$$u_{t,s,a} = \frac{\tau}{2} p_{t,s,a}. \tag{19}$$

Note that we divide $\tau$ by 2 since the gene has an equal chance of moving through a mother or a father. Thus,

$$u_{t,f,1} = \frac{\tau}{2} \cdot 1 = \frac{\tau}{2}, \tag{20}$$

$$u_{t,m,1} = \frac{\tau}{2} \cdot 2w_1 = \tau w_1, \quad u_{t,m,2} = \frac{\tau}{2} \cdot 2w_2 = \tau w_2. \tag{21}$$

Recall that we're tracing a single gene on its journey jumping back in time. To simplify this thought experiment, we've made the assumption that such a gene can only go back to a mother of a single age class and a father belonging to one of two possible age classes. Thus, in each generational step, $p_{f,t,a}$ and $p_{m,t,a}$ are zero for all other age classes, $a$.

Next, define the probability that a random ancestral path at time $t$ is present in an individual of a given age, $a$, and sex, $s$, to be $v_{t,s,a}$. Thus,

$$v_{t,s,a} = \sum_{i=0}^{\infty} u_{t+i,s,a+i} \tag{22}$$

and in particular when $a = 0$,

$$v_{t,s,0} = \sum_{i=0}^{\infty} u_{t+i,s,i}, \tag{23}$$

where we sum over an arbitrary number of years starting from a distant point in the future. Assuming a stable age distribution, the $t$ dependence drops and we obtain the equality

$$v_{f,0} = \sum_{i=0}^{\infty} u_{f,i} = \frac{\tau}{2} \sum_{i=0}^{\infty} p_{f,i} = \frac{\tau}{2} = \frac{\tau}{2} \sum_{i=0}^{\infty} p_{m,i} = \sum_{i=0}^{\infty} u_{m,i} = v_{m,0}. \tag{24}$$

These equations imply that

$$v_{f,0} = u_{f,1} = \frac{1}{2(1+w_2)} = u_{m,1} + u_{m,2} = v_{m,0}, \tag{25}$$

showing that the lifetime reproductive value of newborn females is the same as that of newborn males, as required by Fisher's principle.

We can now determine the total reproductive value over all reproductive age classes given our idealised example. Recall that we consider one reproductive age class for females, $f$, and two for males, $m_1$ and $m_2$.

By definition, when summing over $a \geq 1$ and setting $j = a + i$, we have

$$\sum_a v_{s,a} = \sum_a \sum_{i=0}^{\infty} u_{s,a+i} = \sum_{j=1}^{\infty} \sum_{a=1}^{j} u_{s,j} = \sum_j j u_{s,j} = \sum_a a u_{s,a}. \tag{26}$$

Since we consider only two reproductive age classes, most of these terms are zero and we obtain the following

$$\sum_a v_{f,a} = 1 \cdot u_{f,1} = \frac{\tau}{2} = \frac{1}{2(1+w_2)}, \tag{27}$$

$$\sum_a v_{m,a} = 1 \cdot u_{m,1} + 2 \cdot u_{m,2} = \tau w_1 + 2\tau w_2 = \frac{w_1}{1+w_2} + \frac{2w_2}{1+w_2} = \frac{1+2w_2}{2(1+w_2)}. \tag{28}$$

Note that since, by definition, $w_2 > 0$ whenever males from the older age class win paternities, $\sum_a v_{f,a} < \sum_a v_{m,a}$.

### 3 Conditional probability of male paths

Let $w_1$ and $w_2$ denote the full probabilities that an ancestry path moving through a parent passes through a young father, $m_1$, or an older father, $m_2$, respectively. Since an autosomal gene is equally likely to trace through a mother or a father,

$$P(\text{mother}) = \frac{1}{2}, \qquad w_1 + w_2 = P(\text{father}) = \frac{1}{2}. \tag{29}$$

The quantities $p_{t,m,1}$ and $p_{t,m,2}$ represent the age distribution conditional on the parent being male. Thus,

$$p_{t,m,1} = P(a = 1 \mid s = m) = \frac{P(s = m, a = 1)}{P(s = m)} = \frac{w_1}{w_1 + w_2} = \frac{w_1}{1/2} = 2w_1, \tag{30}$$

and similarly,

$$p_{t,m,2} = P(a = 2 \mid s = m) = \frac{P(s = m, a = 2)}{P(s = m)} = \frac{w_2}{w_1 + w_2} = \frac{w_2}{1/2} = 2w_2. \tag{31}$$

### 4 Stationary distribution of Markov chain

To obtain the stationary distribution, let

$$\pi = (x, y, z)^T. \tag{32}$$

$P$ is a stochastic matrix and $\pi$ is a probability distribution, thus $P\pi = \pi$ and $x + y + z = 1$. Thus, solving the matrix equation and using the fact that $w_2 = 1/2 - w_1$

$$\begin{pmatrix} 0.5 & 0.5 & 0 \\ w_1 & w_1 & 1 \\ w_2 & w_2 & 0 \end{pmatrix} \begin{pmatrix} x \\ y \\ z \end{pmatrix} = \begin{pmatrix} x \\ y \\ z \end{pmatrix}, \tag{33}$$

we get

$$x = \frac{x+y}{2}, \qquad z = w_2(x+y). \tag{34}$$

Thus $x = y$, and $z = 2w_2x$. Normalising gives

$$x + y + z = x + x + 2w_2x = 2x(1+w_2) = 1, \tag{35}$$

so

$$x = y = \frac{1}{2(1+w_2)}, \qquad z = \frac{w_2}{1+w_2}. \tag{36}$$

Therefore,

$$\pi = \left(\frac{1}{2(1+w_2)}, \frac{1}{2(1+w_2)}, \frac{w_2}{1+w_2}\right)^T. \tag{37}$$
